# Regulation of Non-Canonical Proteins Encoded by Small Open Reading Frames via the Nonsense-Mediated Decay Pathway

**DOI:** 10.1101/2023.08.26.554966

**Authors:** Parthiban Periasamy, Craig Joseph, Adrian Campos, Sureka Rajandran, Christopher Batho, James E. Hudson, Haran Sivakumaran, Hitesh Kore, Keshava Datta, Joe Yeong, Harsha Gowda

**Affiliations:** Institute of Molecular and Cell Biology, Agency for Science, Technology, and Research (A*STAR), Singapore 138673, Singapore; QIMR Berghofer Medical Research Institute, Brisbane, QLD, 4006, Australia

**Author notes:** Corresponding authors (Periasamy P), (Yeong J), (Gowda H). **Symbols:** Regeneron Genetics Center, Tarrytown NY. Flow Cytometry Department, Covance Central Laboratory Services, Singapore 609917, Singapore (Sureka Rajandran). Proteomics and Metabolomics Platform, La Trobe University, Melbourne, VIC 3083, Australia.

**Keywords:** UPF1, Nonsense-mediated decay pathway, Mutant transcript, Transcriptional noise, Mutant protein

## Abstract

Immunotherapy interventions relies heavily on neoantigen availability. The human genome encodes non-canonical/mutant proteins that potentially contain neoantigenic peptides. Nevertheless, their typically low expression, potentially moderated by the Nonsense-Mediated Decay (NMD) pathway, restricts their therapeutic utility. In this study, we explored the NMD pathway influence on non-canonical/mutant protein expression, specifically focusing on *UPF1* knockdown. We implemented proteogenomic approaches to ascertain if the encoding transcripts and their respective proteins were upregulated post-knockdown. Complementary to this, we conducted a comprehensive pan-cancer survey of *UPF1* expression and an *in vivo* evaluation of *UPF1* expression in Triple-Negative Breast Cancer (TNBC) tissue. Our empirical results delineated that *UPF1* knockdown precipitates an increase in the transcription of non-canonical/mutant proteins, especially those originating from retained-introns, pseudogenes, long non-coding RNAs, and unannotated biotypes. Furthermore, the analysis revealed that *UPF1* expression was conspicuously high across a range of neoplastic tissues, with protein levels notably amplified in patient derived TNBC tumours in comparison to adjacent tissues. Our study elucidates *UPF1* functional role in attenuating transcriptional noise through the degradation of transcripts encoding non-canonical/mutant proteins. Interestingly, we observed an upregulation of the NMD pathway in cancer, potentially functioning as a “neoantigen masking” mechanism that subdues non-canonical/mutant protein expression. Suppressing this mechanism may unveil a new cadre of neoantigens accessible to the antigen presentation pathway. Our novel findings proffer a solid base for devising therapeutic strategies targeting *UPF1* or the NMD pathway, given the pronounced presence of *UPF1* in malignant cells, thus potentially augmenting immunotherapeutic responses in cancer.

## Introduction

Recently, multiple studies have demonstrated that novel and non-canonical proteins are encoded within non-coding regions of the human genome(1). Most of these newly identified novel proteins are of low molecular weight(2), and the majority are derived from non-coding transcripts with low levels of expression(3). The functions of this growing list of characterized non-canonical proteins(4–6) includes roles in mRNA decapping(7), regulation of calcium ion pump proteins(8, 9), and cancer cell proliferation(10).

Extensive cancer mutome studies have demonstrated that cancer cells contain higher levels of mutations than non-cancerous cells(11). Mutations in the coding regions can result in either loss of function or activation of corresponding proteins. Mutant peptides from these proteins can potentially be presented by major histocompatibility complexes to serve as neoantigens. Additionally, these mutations contribute to the production of defective transcripts(12) with premature stop codons, resulting in truncated or non-canonical protein products. Such mutations may also significantly impair the splicing machinery(13), increasing the diversity of non-canonical transcripts with abnormal open reading frames (ORFs; e.g., mis-spliced transcripts or retained-intron transcripts) and potentially leading to the production of non-canonical proteins(14), which could act as sources of neoantigens. Immunopeptidomics studies(15, 16) have highlighted a substantial population of human leukocyte antigen (HLA)-bound peptides presented on cancer cell surfaces that are derived from non-canonical proteins encoded by long non-coding RNAs (lncRNAs), pseudogenes, untranslated regions (UTRs), and genomically altered reading frames(17). A subset of these peptides is immunoreactive(16), indicating that non-canonical proteins could serve as neoantigens. Despite the successful identification of targetable epitopes for cancer immunotherapy, such non-canonical epitopes are predominantly low in abundance, which has significantly hampered the progress of neoantigen discovery and immunotherapy(18). Furthermore, these epitopes are unique to individual patients(19, 20), thus limiting their application in the personalized medicine setting.

Recently, it has been demonstrated that non-coding RNAs (ncRNAs) and defective transcripts are targeted by a conserved cellular surveillance mechanism known as the nonsense-mediated decay (NMD) pathway(21–23). Downregulation of NMD would be expected to stabilize non-coding and defective transcripts alike, potentially providing an additional source of non-canonical peptides for HLA-bound peptide presentation(14).

In this investigation, our primary objective was to validate the potential role of *UPF1*, a critical protein necessary for NMD, in the regulation of non-canonical proteins encoded by specific transcripts within TNBC cells. To accomplish this, we employed a strategy of *UPF1* knockdown and subsequently assessed the resultant expression levels of these non-canonical proteins. Our experimental findings revealed a notable upregulation of non-canonical proteins, which were encoded by defective and long non-coding transcripts, following the knockdown of *UPF1*. In parallel, we also detected a similar upregulation in unannotated transcripts. Intriguingly, a specific subset of Human Leukocyte Antigen (HLA) transcripts also exhibited a similar pattern of upregulation. These empirical observations led us to postulate that a significant number of these low abundance transcripts and non-canonical proteins may simply represent cellular noise, devoid of any discernible function. Furthermore, we identified that UPF1 is disproportionately abundant in neoplastic cells, rendering it a promising therapeutic target. In summary, our results suggest that the NMD pathway is elevated in cancer cells. This upregulation appears to serve as a mechanism for suppressing the expression of mutant proteins, which could potentially be processed into a source of neoantigens via the antigen presentation pathway. This insight could have significant implications for the development of future cancer therapies.

## Results

### Non-coding transcripts are expressed at low abundance compared to protein-coding transcripts

With the aim of generating a custom protein database for identification of non-canonical proteins, we first conducted transcriptomic profiling of cell lines derived from different breast cancer subtypes (Supplementary Figure 1A). As expected, the median mRNA expression levels across multiple breast cancer cell lines were consistently higher than those of the ncRNA transcripts (*p*-value ≤ 0.0001)(24) (Supplementary Figure 1B and Supplementary File 1). This marked difference in transcript expression led us to further scrutinize the data, in pursuit of potential differences in the expression of ncRNA transcript types that share similar characteristics with protein-coding transcripts, such as pseudogenes. Through density mapping of the log10 transformed Fragments Per Kilobase of transcript per Million (FPKM) values, we discovered a higher degree of overlap between the expression profiles of pseudogene and long non-coding RNA (lncRNA) transcripts as opposed to protein-coding transcripts (refer to Supplementary Figure 1C and Supplementary File 1). These observations lead us to hypothesize that the expression of these non-coding elements is likely regulated by a shared cellular pathway or mechanism.

### Various breast cancer subtypes express novel and non-canonical proteins inconsistently and at low abundance

Given the low abundance of ncRNAs, we hypothesize that encoded novel and non-canonical proteins are also likely less prevalent(2). To identify these proteins in various subtypes of breast cancer cells, we employed an established low molecular weight protein enrichment strategy (Supplementary Figure 2A)(25) that selectively depletes high molecular weight proteins (Supplementary Figure 2B, 2C, 2D and Supplementary File 1). There was a strong correlation between the biological replicates and the proteins in the 10–15 kDa and 15–20 kDa molecular weight ranges within breast cancer subtypes, but a weaker correlation with those in the 0–10 kDa molecular weight range (Supplementary Figure 2E and Supplementary File 1). Utilizing the proteogenomic pipeline shown in Figure 1A, we discovered 2,614 annotated proteins, 344 novel protein isoforms, and 222 non-canonical proteins (Figure 1B and Supplementary File 2). Expression patterns of these proteins were inconsistent and varied across biological replicates and breast cancer subtypes (Figure 1C). We refined the data to confidently identify novel proteins based on certain criteria, such as consistent detection across replicates, a minimum of two peptide spectrum matches, and identification in at least two breast cancer cell lines. This led to the identification of 39 novel proteins unique to triple-negative breast cancer (TNBC) cell lines, as annotated in the OpenProt(26, 27) protein database (Figure 1D and Supplementary File 2). One such protein, IP_156279, was detected with a total peptide spectrum match count of 7. We also observed differential expression of these novel and non-canonical proteins between TNBC and non-TNBC cell lines (Figure 1E and Supplementary File 2). Interestingly, a long non-coding RNA (lncRNA)-encoded protein (IP_786419) was found in eight breast cancer cell lines, with higher expression in TNBC subtypes. Predictive modeling using DeepLoc(28) suggested this protein is localized within mitochondria.

**Figure 1.**
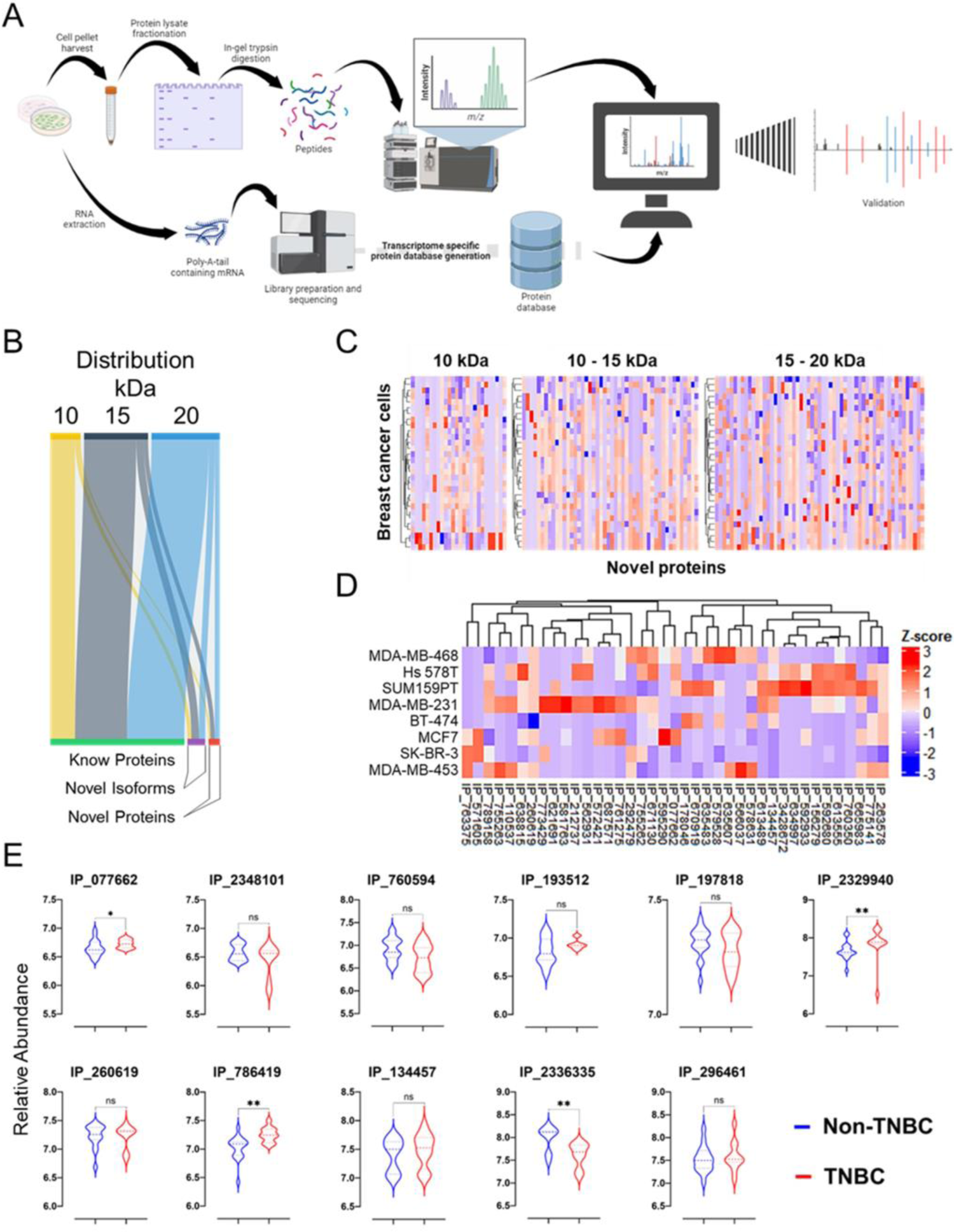
Discovery of novel proteins in breast cancer cell lines. (**A**) Workflow for novel protein discovery using a proteogenomics pipeline. Paired proteomics and transcriptomics data were generated, integrated, and analyzed. Newly discovered novel protein peptide spectrum matches (PSMs) were validated in silico using Prosit and/or synthetic peptides. (**B**) Distribution plot of total proteins identified from all breast cancer cell lines in this study. Identified proteins were categorized as known proteins, novel isoforms, or novel proteins. (**C**) Heatmap of Z-score transformed PSM counts of all novel proteins identified. (**D**) Heatmap of Z-score transformed PSM counts of high confidence novel protein identified. (**E**) Detection and quantification of representative differentially modulated novel proteins in triple negative and non-triple negative breast cancer cell lines using log10 transformed normalized abundance values. Statistical analysis was performed using an unpaired t-test (parametric test). Three independent biological experiments (n = 3) were performed. ns – not significant, * p ≤0.05 and ** p ≤ 0.01.

### LncRNA-HELLPAR encodes a novel protein in TNBC that is rapidly degraded

Our proteogenomic analyses identified a novel protein encoded by the lncRNA-HELLPAR(29, 30) transcript in TNBC cell lines; this lncRNA maps to Chr12 (q23.2) (Supplementary Figure 3A). Using one-step RT-PCR (Supplementary Figure 3B and Supplementary File 2), we confirmed varying levels of lncRNA-HELLPAR transcript expression across all eight breast cancer cell lines (Supplementary Figure 3B). Unsurprisingly, RNA-seq revealed that the abundance of this transcript was relatively low (Supplementary Figure 3C), similar to the previously published expression profiles of known lncRNAs(24, 31, 32) as well as our own RNA-seq data (Supplementary Figure 1C). LncRNA-HELLPAR expression was highest in the TNBC cell lines SUM159PT and Hs 578T. The HELLPAR transcript encodes a 61-amino acid protein with a predicted N-terminal glycosylation site (Supplementary Figure 3D). The HELLPAR protein (IP_760350) was detected at high abundance in TNBC cell lines Hs 578T and SUM159PT (Figure 1D) and has been detected in a previous study(25). Using a synthetic peptide to confirm the spectral assignment (Supplementary Figure 3D and Supplementary File 2), we then validated the presence and quantity of this peptide across a panel of cell lines comprising all breast cancer subtypes. Targeted proteomic analysis confirmed abundant expression of HELLPAR in the TNBC cell lines MDA-MB-231, Hs 578T, and MDA-MB-468 compared to other breast cancer subtypes (Supplementary Figure 3E and Supplementary File 2). MS/MS searches of a publicly available high-resolution, deep fractionated human proteome dataset consisting of different tissues(33) revealed that the HELLPAR-encoded protein was expressed in multiple adult and fetal tissues (Supplementary Figure 3F and Supplementary File 2).

In an effort to decipher the functionality of the protein encoded by HELLPAR, we prompted its overexpression in SUM159PT cells, as demonstrated in Supplementary Figure 3G. However, our attempts to detect the HA-GFP-tagged HELLPAR protein, using a plasmid-based overexpression system, proved ineffective (Supplementary Figure 3H). This was despite the detection of minimal levels of the transcript in transfected cells, as evidenced by one-step RT-PCR (Supplementary Figure 3I). As a strategy to counteract this low abundance, we engineered a HA-GFP-tagged HELLPAR modified mRNA and transfected it into the SUM159PT breast cancer cells, thereby facilitating transient overexpression of HELLPAR (Supplementary Figure 3J). We found that transfection with the HELLPAR novel ORF linked to GFP-HA (devoid of a methionine start codon) at the C-terminus yielded expression of the HELLPAR-GFP-HA protein (Supplementary Figure 3K). This strongly indicates that the protein translation was propelled by the HELLPAR ORF. Yet, in spite of the incorporation of a robust protein translation initiation sequence, the Kozak sequence, at the onset of the ORF, HELLPAR expression was still subpar when compared to the GFP-HA transfection control (Supplementary Figure 3L). This observation leads us to hypothesize that the HELLPAR transcript or protein may be subject to swift degradation post-synthesis.

### The NMD pathway modulates levels of transcript encoding novel and non-canonical proteins

A large number of non-canonical proteins, including the lncRNA-HELLPAR-encoded protein, along with their respective transcripts, exhibit low and inconsistent expression profiles. This observation leads us, in line with previous studies(34), to posit that many such transcripts might essentially constitute transcriptional noise. It is plausible that the abundance of these transcripts is controlled via an evolutionarily preserved mechanism, such as the NMD pathway(23, 35). In order to scrutinize if the perturbation of the NMD pathway has any influence on the production of non-canonical proteins encoded by ncRNAs, we executed a siRNA-mediated knockdown of UPF1, a crucial component of the NMD pathway. This was performed in TNBC cell lines MDA-MB-231 and SUM159PT. Following this, we undertook a comprehensive analysis of the transcriptomic and proteomic profiles in alignment with the workflow delineated in Figure 1A. We confirmed the efficacy of UPF1 knockdown at both RNA and protein levels in these cell models (Figure 2A and 2B). Transcriptomic analysis revealed that most protein-coding transcripts remained unchanged following UPF1 knockdown, while ncRNA and NMD transcripts showed an upregulation. This pattern was consistent across both breast cancer cell lines analyzed (Figure 2C and Supplementary File 3), indicating that our NMD model functioned as expected and concurred with previous findings (Supplementary Figure 4A, 4B and Supplementary File 3)(31).

**Figure 2.**
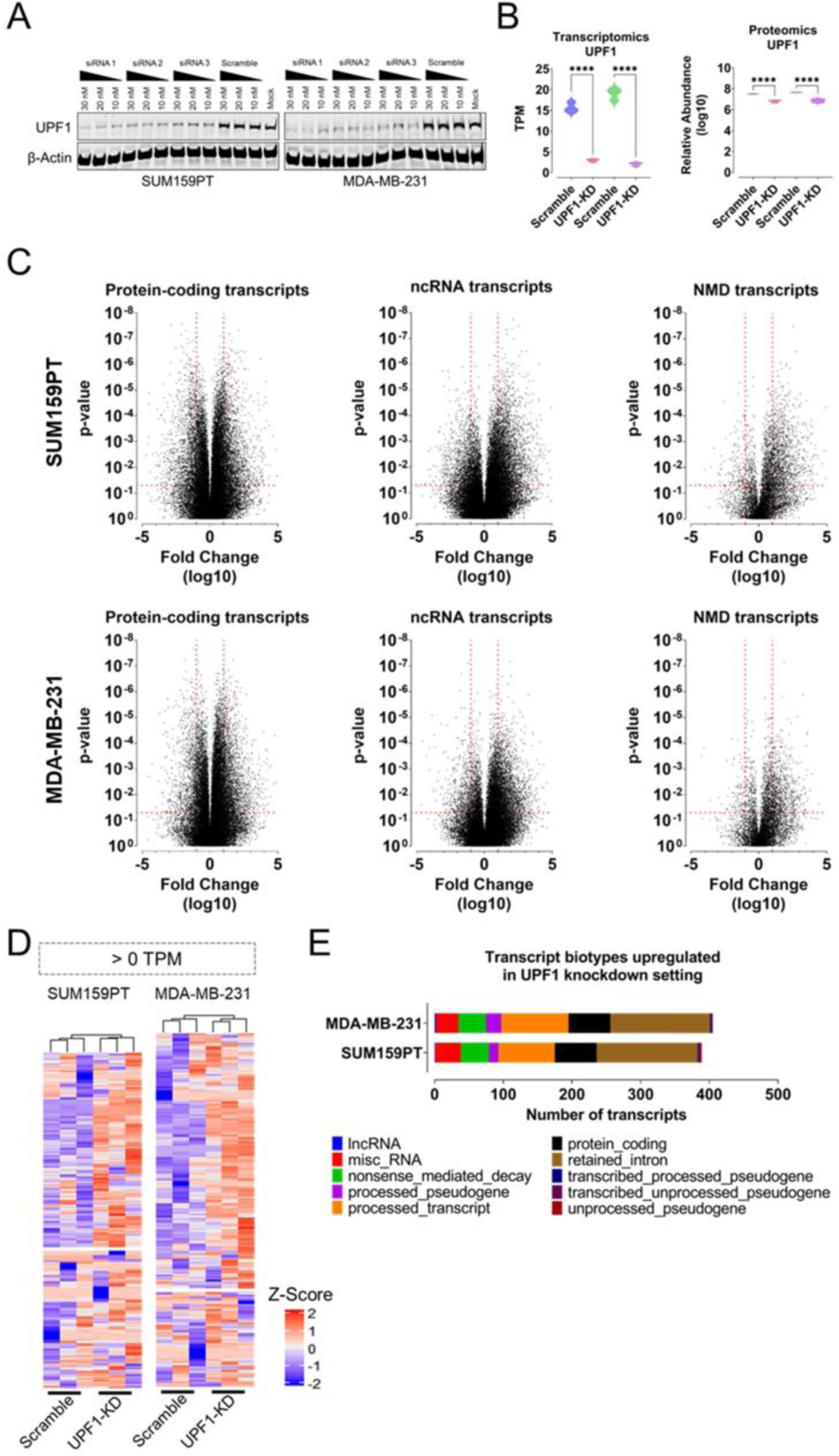
ncRNAs and mis-spliced transcripts are targeted by the nonsense-mediated decay pathway. UPF1 siRNA knockdown in the breast cancer cell lines SUM159PT and MDA-MB-231 and subsequent proteogenomic characterization. Three days after siRNA treatment, total RNA and proteins were extracted as per supplementary Figures 1 and 2, respectively, and processed. (**A**) Validation of various UPF1 siRNAs transfected at different concentrations into the breast cancer cell lines SUM159PT and MDA-MB-231. Western blotting was performed using beta-actin as a loading control. (**B**) Transcriptomic and proteomic validation of siRNA-mediated knockdown of UPF1 in the breast cancer cell lines SUM159PT and MDA-MB-231. UPF1 RNA and protein levels were plotted to determine knockdown efficiency. (**C**) Transcriptomic profiles of UPF1 targeting siRNA-treated SUM159PT and MDA-MB-231 cells. Log10 fold change values for differentially modulated protein-coding, ncRNA, and NMD transcripts were plotted; the corresponding p-values were determined using two tailed t-tests. (**D**) Heatmap showing Z-scores (transformed log10 normalized protein abundance values converted to Z-score) of differentially regulated novel isoforms and novel proteins in cells transfected with scrambled or UPF1 knockdown siRNAs. (**E**) Transcript biotypes of upregulated novel isoforms and novel proteins following UPF1 knockdown. Transcript biotype is based on GenCodeV35 classification. **** p ≤ 0.0001.

Having demonstrated the functionality of our NMD model, we then integrated our transcriptomic and proteomic data into our proteogenomic pipeline, we observed an upregulation of numerous novel proteins in both *UPF1* knockdown cell lines (Figure 2D and Supplementary File 3). Notably, more non-canonical proteins derived from low abundance transcripts (<1 TPM) showed an increase compared to high abundance transcripts (>1 TPM). Most novel proteins upregulated were encoded by retained-intron transcripts, typically degraded by the NMD pathway (Figure 2E)(36).

However, not all ncRNAs exhibited upregulation with UPF1 knockdown. Certain well-known lncRNAs like ENOX, CHROME, TARS1-DT, PPP1R21-DT, and MIR762HG maintained stable expression levels (Supplementary Figure 4E-I). Additionally, specific novel proteins encoded by ncRNAs remained unaffected by NMD pathway modulation, possibly suggesting authentic mRNA transcription and translation for functional protein production. Examples include IP_591189, IP_665356, IP_637007, IP_107612 and IP_734753 (Supplementary Figure 4J-O).

### *UPF1* knockdown results in upregulation of HLA and novel transcripts

Given the successful elevation of cellular abundance of ncRNA-derived novel and non-canonical proteins, we postulated an upregulated sampling and presentation of these proteins in UPF1 knockdown cell lines. Our examination of RNA-seq data for HLA transcript expression disclosed a consistent upregulation of certain HLA transcripts in UPF1-knocked-down MDA-MB-231 and SUM159PT breast cancer cell lines (Figure 3A).

**Figure 3.**
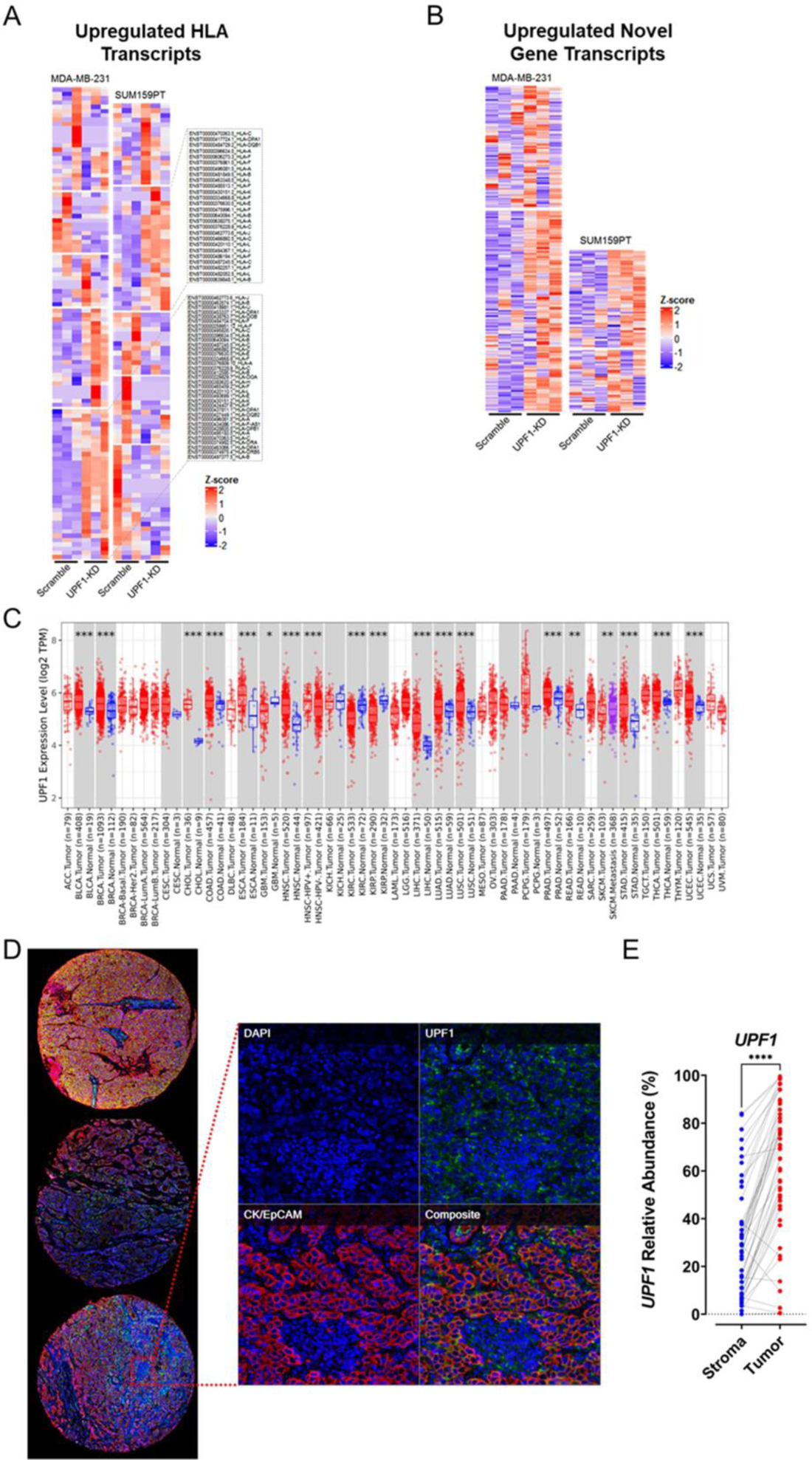
Perturbation of the nonsense-mediated decay pathway via siRNA-mediated UPF1 upregulates HLA/novel transcripts and UPF1 is highly expressed in vivo in various types of cancer, while being expressed at low levels in healthy tissues. SUM159PT and MDA-MB-231 cells were processed as per Supplementary Figure 1A (transcriptomics). (**A**) Detected transcripts were filtered for HLA transcripts, and Z-score transformed FPKM values were plotted. (**B**) Additionally, pre-processed raw fastq files were realigned using HISAT2, and de novo assembly of novel transcripts currently unannotated in the GeneCodeV35 GTF files was performed using StringTie. After filtering for novel transcripts, Z-score transformed FPKM values were plotted. (**C**) Expression level of UPF1 RNA in both healthy and tumor tissues, represented in blue and red, respectively. (**D, E**) Tumor tissue sections derived from patients with TNBC (n = 48). The sections were stained with antibodies against UPF1, CK, and EpCAM, and images were captured. Subsequently, the abundance (%) of UPF1 was quantified in the tumor and stroma regions and plotted. Two-tailed t-tests were performed to determine the corresponding p-values. **** p ≤ 0.0001.

Most transcription initiation events mediated by RNA polymerase II in eukaryotes have been proposed to constitute transcriptional noise, resulting in biologically insignificant RNA(34). We hypothesized that alongside the upregulation of ncRNA transcription within the cellular milieu, novel transcripts not yet annotated in the GENCODE database would also manifest in high abundance in the UPF1 knockdown scenario. Addressing this conjecture, we reanalyzed our transcriptomic data via de novo assembly of novel transcripts. Intriguingly, we found a consistent upregulation of numerous novel transcripts in the UPF1 knockdown context across three biological replicates and two distinct breast cancer cell lines (Figure 3B). This suggests a role for UPF1 in mitigating transcriptional noise.

### *UPF1* exhibits high RNA and protein expression in multiple cancer types and patient-derived triple-negative breast cancer tissue, while expression is low in normal tissue

Our research to date has indicated that siRNA mediated UPF1 knockdown can enhance the expression of novel and non-canonical proteins derived from ncRNAs and defective transcripts. These proteins may act as neoantigens, potentially amplifying immunotherapy responses in cancer. To assess the applicability of our approach in improving immunotherapy responses across diverse cancer types, we analysed UPF1 expression levels in various tumor types and their corresponding healthy tissues. A transcriptomic analysis of data from The Cancer Genome Atlas and The Genotype-Tissue Expression project revealed a significant upregulation of UPF1 in most cancer subtypes, compared to their respective normal tissues (Figure 3C). Immunohistochemical staining of UPF1 and CK/EpCAM (a tumor cell marker) in 48 patients derived TNBC sections corroborated the transcriptomic data, demonstrating a higher expression of UPF1 in tumor cells compared to stromal cells (Figure 3D and 3E). Collectively, our findings suggest an upregulated NMD pathway in cancer cells, potentially acting as a suppressor of mutant protein expression. These suppressed proteins, upon processing via the antigen presentation pathway, could potentially serve as a source of neoantigens.

## Discussion

In this study, we investigated the ability of the NMD pathway to modulate the expression of transcripts that encode non-canonical or mutant proteins. To address this aim, we mapped the low molecular weight proteome using breast cancer lines as a model. We then investigated the potential of increased expression of novel and non-canonical proteins derived from ncRNA and defective transcripts to expand the source of neoantigens as an approach to augmenting immunotherapy responses in cancer.

First, our RNA-seq data indicated that non-coding transcripts (including defective transcripts) were expressed at significantly lower levels than protein-coding transcripts. Fascinatingly, the transcripts produced by pseudogenes (which resemble functional genes) were expressed at similarly low levels, despite containing a poly-A tail, which plays a pivotal role in RNA stability(37). Furthermore, we obtained analogous results by analyzing an independent, publicly available dataset (based on the HeLa cancer cell line)(31). We speculated that these non-coding and defective transcripts exhibited shorter half-lives than protein-coding transcripts and moreover, that degradation of these transcripts may occur via a conserved cellular pathway or mechanism.

Second, using our proteogenomic approach, we identified several novel proteins encoded by ncRNA and defective transcripts. Interestingly, most of the novel proteins displayed inconsistent expression profiles within biological replicates and among breast cancer subtypes, possibly due to aberrant transcription and/or translation. Therefore, we applied additional stringent filtering criteria to validate the authenticity of these novel proteins and shortened the list to 39, including proteins derived from pseudogenes, lncRNAs, and UTRs. Despite identifying multiple novel proteins, the corresponding peptide spectra were of poor quality and unreliable. Nonetheless, we were able to identify a lncRNA-encoded protein (accession ID IP_786419) that was predicted(28) to be localized to mitochondria and was expressed in all the breast cancer cell lines tested. Additionally, we functionally characterized the protein encoded by lncRNA-HELLPAR, which had been characterized at the RNA level to have a role in trophoblast invasion and maternal health(29, 30), and exhibited both nuclear and cytoplasmic localization. Analysis of global proteomic profiling data from human adult and fetal tissues(33) revealed HELLPAR-encoded protein expression in multiple tissues. However, we were unable to achieve stable overexpression of the HELLPAR-encoded protein using either DNA- or RNA-based systems. The low abundance and inconsistent expression of most of these novel/non-canonical proteins, including the lncRNA-HELLPAR-encoded protein, prompted us to evaluate whether such proteins do indeed have physiological functions or are simply the result of transcriptional and translational noise detected by the more sensitive methods currently in use. It should be noted that most novel and non-canonical proteins identified by RNA-seq and/or proteomics are poorly conserved across species(1, 10), while the lncRNAs that encode these proteins are often expressed at extremely low levels(3). This issue is exemplified in our overexpression study of the lncRNA-HELLPAR transcript and encoding protein. All of this evidence prompted us to speculate whether these transcripts and proteins, including lncRNA-HELLPAR and the encoded protein, may simply be transcriptional(32, 34) and translational noise(38) respectively, without apparent function. If so, a cellular mechanism is required to keep such transcriptional/translational noise in check. In addition, considering the widespread transcription(34) in most eukaryotic genomes, we hypothesized that a RNA surveillance mechanism to degrade these transcripts should be conserved across species. The NMD pathway, a highly conserved RNA surveillance mechanism, targets transcripts with premature stop codons for degradation (23). This process is activated by persistent exon-junction-complex (EJC) attachment during translation. The EJC, bound to exon-exon junctions of multi-exonic transcripts post-splicing in the nucleus, is subsequently relocated to the cytoplasm, where it interacts with ribosomes. As protein synthesis proceeds, ribosome scanning dislodges the EJC from these transcripts. However, an exon harboring a premature stop codon, barring the terminal exon, prevents EJC dislodgement downstream, consequently triggering the NMD pathway to degrade the transcript. The NMD pathway role extends beyond targeting mRNAs with premature stop codons, potentially encompassing numerous ncRNAs and defective transcripts encoded by the human genome(39). We hypothesized that such transcripts, often bound non-specifically by ribosomes in the cytoplasm (40), might be targeted by the NMD pathway post-translation, contributing to the scarcity of most sORF-encoded proteins(23, 39). This theory was evaluated using transcriptomic and proteomic profiling of TNBC cell lines post-UPF1 knockdown, a pivotal NMD pathway component. The ensuing ncRNA and novel protein overexpression suggests the NMD pathway modulates noise from many sORF-encoded proteins, which likely lack function. Investigations into the functionality of lncRNA- and sORF-encoded proteins are ongoing. However, our findings underscore the necessity for caution when studying low abundance proteins, as their detectability may not equate to physiological relevance.

Intriguingly, our experimental outcomes suggest that the inhibition of NMD could potentially lead to an augmented expression of non-conventional, *de novo*, and mutant proteins within the cellular milieu. Crucially, given the rapid degradation of these proteins, they are poised for presentation by Major Histocompatibility Complex (MHC) molecules on the surface of neoplastic cells(16). This mechanistic pathway could effectively enhance the neoantigenic load presented by malignant cells, thereby augmenting the effectiveness of checkpoint blockade therapeutic approaches (Figure 4). Moreover, considering the ubiquitous expression of certain non-conventional proteins, the inhibition of NMD could generate universal neoantigens that could be targeted across a diverse array of neoplastic pathologies. We are presently conducting preclinical investigations utilizing immunocompetent murine models to verify the practicality of this therapeutic strategy. Collectively, our data propose that a combinatory therapeutic regimen encompassing both NMD inhibitors and checkpoint inhibitors may broaden the applicability of immunotherapy to a larger patient population without necessitating personalized neoantigen identification.

**Figure 4.**
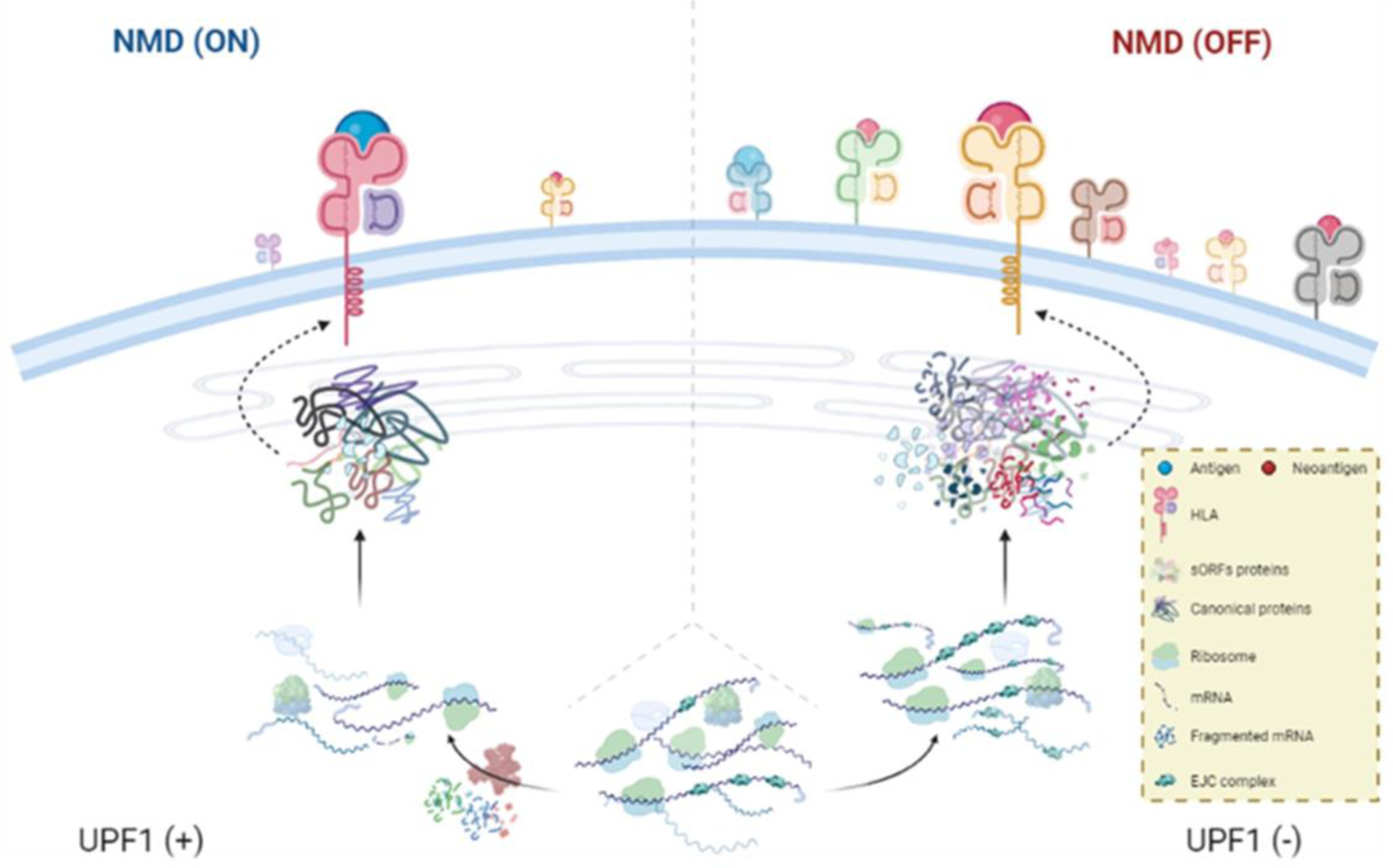
Model of neoantigen/non-canonical peptide presentation in the presence or absence of nonsense-mediated decay. In the presence of nonsense-mediated decay, mis-spliced, ncRNA and/or aberrant transcripts containing premature stop codons are subjected to decay as a result of exon-junction-complex (EJC) retention. Only authentic protein-coding transcripts are stably retained within the cellular environment. In the absence of nonsense-mediated decay, aberrant transcripts are retained within the cellular environment and translated along with authentic protein-coding transcripts. This results in increased cellular noise in terms of the transcriptome/proteome, as well as sampling of aberrant translation protein products that, along with non-canonical proteins, are processed to shorter 8–14-mer peptides with the potential to act as an enriched source of neoantigens for HLA presentation.

Immunotherapy, particularly checkpoint blockade therapy, has transformed cancer treatment, notably enhancing response rates in metastatic melanoma and lung cancers. However, its efficacy can be limited by the low neoantigen burden on cancer cells. Recent studies suggest that non-traditional, novel, and mutant proteins encoded in the human genome may serve as a robust source of cancer neoantigens. Amplifying the mutant protein load intracellularly and increasing mutant peptide presentation, facilitated by NMD inhibitors under investigation, could provide a targeted approach to tumor destruction without impacting healthy cells.

## Materials and methods

### Cell lines

MDA-MB-467, Hs 578T, SUM-159-PT, MDA-MB-231, BT-474, MCF7, SK-BR-3 and MDA-MB-453 were obtained from the American Type Culture Collection (ATCC) and cultured and maintained as per ATCC recommendations. All cell lines were authenticated by short tandem repeat profiling and tested for mycoplasma.

### RNA extraction

Breast cancer cells were seeded in 6-well plates at 5 × 10^4^ cells per well and cultured as per ATCC requirements to 80% confluence. The culture supernatants were removed, and the cell monolayers were washed three times in ice-cold phosphate-buffered saline (PBS). Subsequently, total RNA was extracted using 500 μL TRIzol™ Reagent (Cat.# 15596018 - CA, USA) according to the manufacturer’s instructions. The RNA pellets were then resuspended in 30 μL ice-cold nuclease-free water and stored at −80°C prior to use.

### RNA-seq data analysis

Raw FASTQ files were processed using Trimmomatic V0.36 (ILLUMINACLIP, SLIDINGWINDOW 4:15 and MINLEN:36 functions) and the quality was evaluated using FastQC V0.11.9. For alignment using Salmon(41), the GENCODE human release V35 transcriptome FASTA file was first used to build an index set using the default Salmon index function with k-mers of length 31. Using the generated index, read-mapping was performed using the Salmon quant function with the parameters auto library type (-l A) and mimic strict bowtie2 alignment (--mimicStrictBT2). Output alignment files were merged into a non-redundant matrix file for further downstream data analysis.

For the discovery of novel transcripts (*de novo* mode), processed raw FASTQ files were aligned using HISAT2(42) V2.0.4. First, human genome indexes were built using the HISAT2 build function with the GENCODE V35 GRCh38 primary assembled genome FASTA file. Second, the GENECODE V35 GRCh38 comprehensive gene annotation file was used to build the known splice site index file. Thereafter, the HISAT2 alignment function was used with reverse-forward library type (--rf) and known splice site infile (--know-splicesite-infile) as parameters. The alignment output SAM file was converted into a coordinate-sorted BAM file using the Samtools V1.9 view and sort functions sequentially. Read assembly was performed with the library type reverse-forward (--rf) function of StringTie V1.3.5 and the output GTF files were merged to generate a global unified set of transcripts (isoforms) across multiple RNA-seq samples with the following functions: minimum transcript length of 200 nt (-m 200), keep transcripts with retained introns (-i), and minimum input transcript fragments per kilobase million (FPKM) of 0.000001 (-F 0.000001). The read assembly was then repeated using this file and the GTF files were merged using an in-house Python script to generate a matrix file for further downstream data analysis.

For novel transcript heatmap plotting, we used R-package ComplexHeatmap version 2.6.6 with Z-score transformed log10 normalized TPM values. To generate custom cell line-specific protein databases, protein sequences were extracted from the OpenProt(27) full protein database using Ensembl identifiers with an in-house Python script. Thereafter, a list of common contaminants was appended to the custom protein database.

### One-step lysis and high molecular weight protein depletion

Breast cancer cells were seeded in two 15-cm dishes at 5 × 10^6^ cells per well and cultured as per ATCC requirements to 80% confluence. The culture supernatants were removed, and the cell monolayers were washed three times in ice-cold PBS. The cells were then trypsinized and pelleted in 15-mL protease-free tubes for storage at −80°C. For high molecular weight protein depletion, cell pellets were processed according to a previously published protocol(25).

### In-gel peptide preparation for LC-MS/MS

For in-gel digestion, Coomassie-stained protein bands obtained after SDS-PAGE (0–10 kDa, 10–15 kDa and 15–20 kDa; each kDa range gel band was processed individually) were destained using 40 mM TEABC buffer (Sigma – Cat. #T7408-500ML) in 40% acetonitrile. Subsequently, the proteins were reduced using 5 mM dithiothreitol (final concentration) with gentle shaking at 60°C for 30 min. Next, the proteins were alkylated by incubation with 20 mM iodoacetamide (final concentration) in the dark at room temperature for 10 min. Reduced and alkylated samples were digested in-gel using 200 μL (or a sufficient volume to cover all the gel pieces) of MS grade trypsin (Promega – Cat.#V5111), diluted to 10 ng/µL in 40 mM TEABC buffer in 1.5-mL proteomics-compatible tubes. After 16 h, the trypsin was inactivated by adding 0.1% formic acid (final concentration) (Pierce^TM^ – Cat.#85178) and shaking vigorously at 37°C for 10 min. The supernatants were collected, and the digested peptides were extracted in 300 μL 5% formic acid/40% acetonitrile, twice with vigorous shaking at 37°C for 15 min. A final peptide extraction was performed using 500 μL 100% acetonitrile with vigorous shaking at 37°C for 20 min or until the gel became opaque. All collected supernatants were pooled and vacuum-dried in a GeneVac EZ2 using the DriPure® 30°C setting. Dried peptide pellets were resuspended in 200 μL of 0.1% formic acid, and peptide clean-up was performed using in-house prepared C18 stage tips (3M™ Empore™ – Cat.#14-386-2). Cleaned-up peptides were vacuum-dried in a GeneVac EZ-2and stored at −20°C.

### siRNA mediated *UPF1* knockdown

The cancer cell lines SUM159PT and MDA-MB-231 were seeded in 6-well tissue culture plates at 2.5 ×10^5^ cells per well and cultured for approximately 24 h to 60% confluence. Subsequently, the cell monolayers were washed twice with sterile tissue culture grade PBS, and 750 μL additive-free DMEM was added to each well. Cells were transfected with 10 nM siRNA targeting *UPF1* (5’-GTGACGAGTTTAAATCACAAATCGA-3’, 5’-CAGCATCTTA TTCTGGGTAATAAAA-3’, or 5’-TCAAGGTCCCTGATAATTATGGCGA-3’) or scrambled siRNA (non-targeting control) using 6 μL Lipofectamine™ RNAiMAX (Invitrogen™ - Cat.# 13778075). After 24 h, culture media were removed, and the cells were maintained as per ATCC recommendations. At 60 h post-transfection, the cells were washed once with PBS and processed for low molecular weight protein enrichment as described previously(25) or for total RNA extraction.

### Mass spectrometry analysis

Peptides were resuspended in 30 μL 0.1% formic acid, and the concentration was estimated using the NanoDrop™ One spectrophotometer (ɛ205 = 31 method). Peptides (approximately 2 μg) were analyzed using the Thermo Scientific Orbitrap Fusion Tribrid mass spectrometer interfaced with the Waters nanoACQUITY UPLC system. Peptides were loaded onto a Waters Acquity C18 UPLC trap column (180 μm ID × 2 cm, 5 μm particle size, 100-Å pore size) using 5% solvent B (0.1% formic acid in acetonitrile) at a flow rate of 5 μL/min for 6 min. Peptides were then resolved on a Waters nanoACQUITY BEH C18 analytical column (75 μm ID × 20 cm, 1.7 μm particle size, 170 Å pore size) using a flow rate of 300 nL/min. The elution gradient used was 12%–40% solvent B, followed by a ramp up to 95% solvent B for 5 min, with a total run time of 60 min. Solvent solutions were prepared using Optima LC-MS grade acetonitrile, water, and formic acid (Thermo Scientific). The mass spectrometer was operated in positive mode for data-dependent acquisition. Full-scan MS1 spectra were acquired at a resolution of 60,000 from 350–1800 m/z with a 3 s cycle time and an automatic gain control target value of 1 × 10^6^. MS/MS spectra were acquired at a resolution of 15,000 with an isolation window of 1.2 and an automatic gain control target of 5 × 10^4^. Additional liquid chromatography and mass spectrometry method details can be obtained from the proteomics RAW files.

### Proteomics data analysis

MS/MS spectra were searched using the FragPipe (version 16.0) interface coupled with MSFragger(43) search engine (version 3.3) and Philosopher(44) data analysis software (version 4.0). FragPipe default label-free quantification with match-between-run workflow was performed using the UniProt canonical human proteome database (downloaded on 1^st^ December 2021 and appended with a list of common contaminants), changing the match-between-runs MS1 ion retention time tolerance window to 3 min.

For novel protein discovery and quantification, MS/MS spectra were searched as described above, using either the OpenProt(27) full proteome database (downloaded on 1^st^ December 2021 and appended with a list of common contaminants) or a custom 3-frame translated cell line-specific database generated using RNA-seq data. Output MSstats files were processed using R-package MSstats(45) version 3.22.1, Stringr version 1.4.0, and readr version 2.1.1 without imputations. Detected and quantified proteins were log10-transformed, and group comparisons were performed. Differentially expressed proteins (DEPs) between TBNC and non-TNBC cell lines were analyzed using *t*-tests (unpaired, parametric test, two-tailed) in GraphPad Prism version 9.3.1.471. Heatmaps were plotted using R-package ComplexHeatmap version 2.6.6 with Z-score transformed log10 normalized abundance values.

### Multiplex immunohistochemistry

To perform multiplex immunohistochemistry (mIHC) staining, 4-µm thick formalin-fixed paraffin embedded (FFPE) tissue microarray (TMA) sections of TNBC were used. The Leica Bond Max autostainer (Leica Biosystems, Melbourne, Australia), Bond Refine Detection Kit (Leica Biosystems, Newcastle Upon Tyne, UK), and Opal 6-Plex Detection Kit for Whole Slide Imaging (Akoya Biosciences, Marlborough, MA, USA) were employed for the staining process. The FFPE tissue sections were deparaffinized, rehydrated, and subjected to repeated cycles of heat-induced epitope retrieval, followed by incubation with primary antibodies (UPF1 - cataloge number: ab109363, CK-cataloge number: ab93741 and EpCAM – cataloge number: ab223582) and secondary antibodies (Bond Refine Kit), and Opal tyramide signal amplification (TSA) (Akoya Biosciences, Marlborough, MA, USA). After all markers were stained, spectral DAPI (Akoya Biosciences, Marlborough, MA, USA) was used to counterstain the nuclei. Finally, the slides were mounted with ProLong Diamond Anti-fade Mountant (Molecular Probes, Life Technologies, USA) and developed in the dark at room temperature for 24 h. The images were captured using Zeiss AxioScan 7 (Carl Zeiss, Germany) and analyzed and scored using HALOTM (Indica Labs).

## Authors’ contributions

Conceptualization, PP, and HG; methodology, PP, CJ, AC, SR, CB, HS, and HK; formal analysis, PP, SR, CB, JH, KD, HS, HK, and HG; investigation, PP; resources, JH, JY, and HG; writing – original draft, PP; writing – review & editing, PP and HG; supervision, JH, JY, and HG; funding acquisition, JH, JY and HG.

## Acknowledgements

We would like to express our gratitude to several parties for their valuable contributions to this work. Firstly, we would like to thank Prof. Khanna from QIMRB for providing breast cancer cell lines, and QIMRB Scientific Services for assisting with mycoplasma testing and cell line authentication. Additionally, we would like to acknowledge the excellent administrative and technical support provided by the QIMRB sequencing facility and high-performance computing department. Funding: The mIHC work was funded by a grant from the A*STAR BIOMEDICAL ENGINEERING PROGRAMME (Project No: C211318003) of the Singapore National Medical Research Council (MOH-000323-00, OFYIRG19may-0007) and IAF-PP (HBMS Domain): H19/01/a0/024-SInGapore ImmuNogrAm for Immuno-Oncology (SIGNAL) awarded to J.Y. Finally, we would like to extend our gratitude to H.G. NHMRC funding. H.G. is a NHMRC R.D. Wright Fellow.

## Supplementary material

**Supplementary figure 1.**
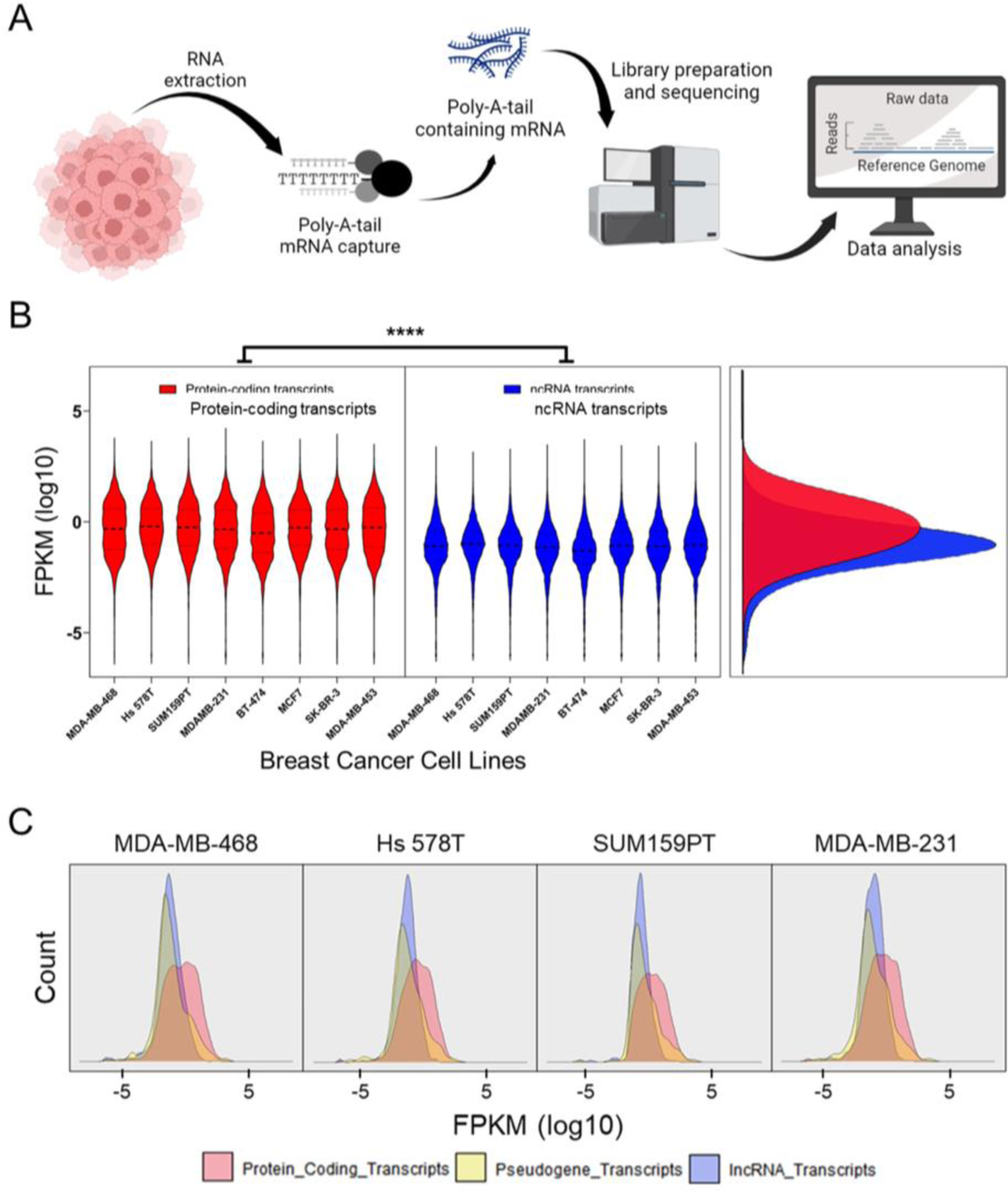
Higher abundance of protein-coding transcripts compared with non-coding transcripts across a panel of breast cancer cell lines. (**A**) Workflow for transcriptomics. (**B**) Comparison of protein-coding and non-coding poly-A tail-containing transcript abundance across various breast cancer cell lines. Nested t-tests were performed to compare the log10 transformed FPKM of protein-coding and non-coding transcripts; **** p ≤ 0.0001. (**C**) Comparison of transcript subtype-specific abundance (protein-coding, pseudogene, and lncRNA transcripts) within the sample cell lines MDA-MB-468, Hs 578T, SUM159PT, and MDA-MB-231.

**Supplementary figure 2.**
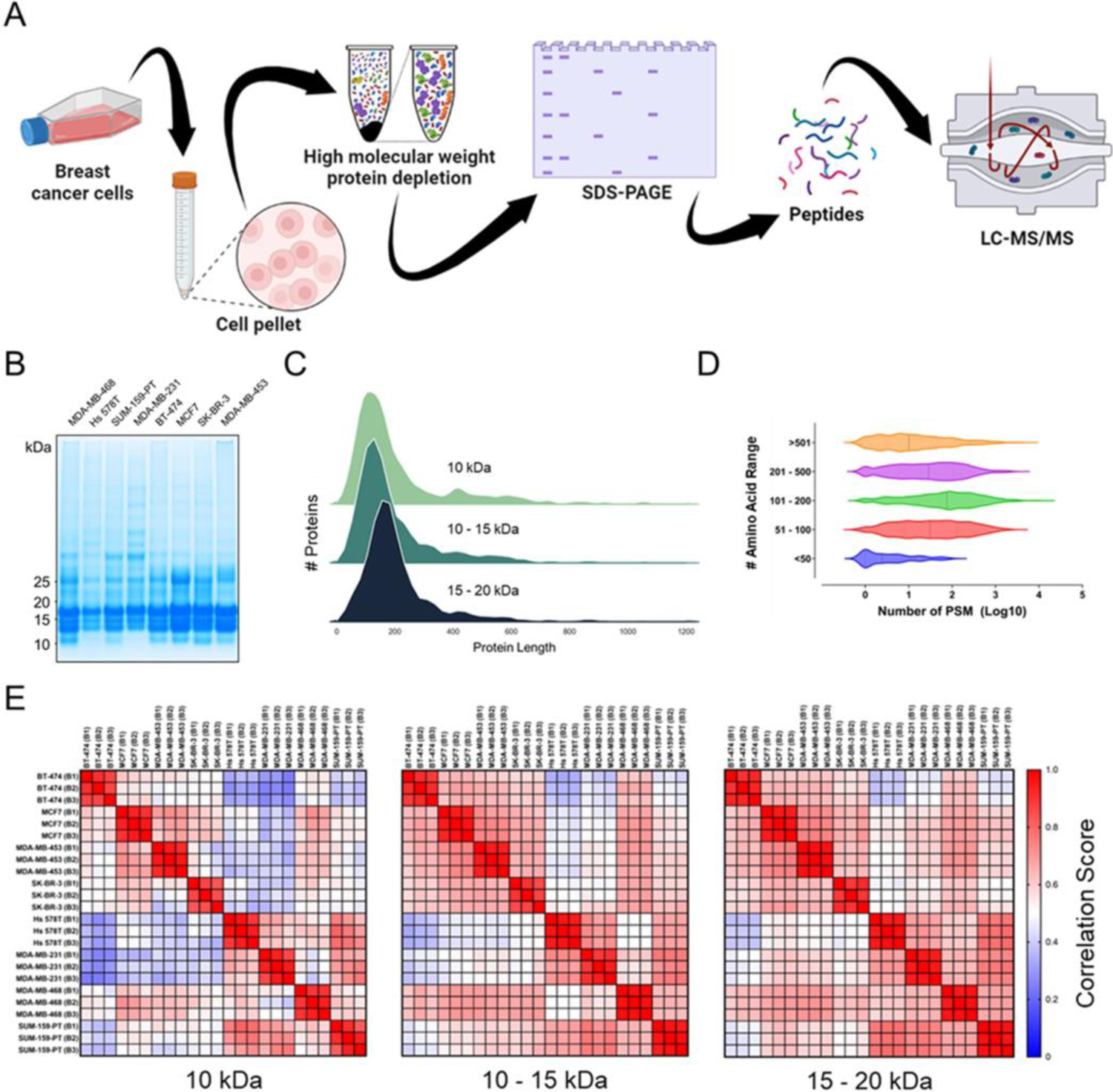
Small protein enrichment in breast cancer cell lines. (**A**) Workflow for small protein enrichment. (**B**) Representative gel showing small protein enrichment in a panel of breast cancer cell lines. (**C**) Density plot of total identified proteins from small protein enrichment workflow. (**D**) Total log10 normalized peptide spectral match (PSM) count of identified peptides from all cell lines for the three molecular weight ranges. (**E**) Correlation plot of detected and quantified proteins in different breast cancer cell lines (n = 3) using abundance values.

**Supplementary figure 3.**
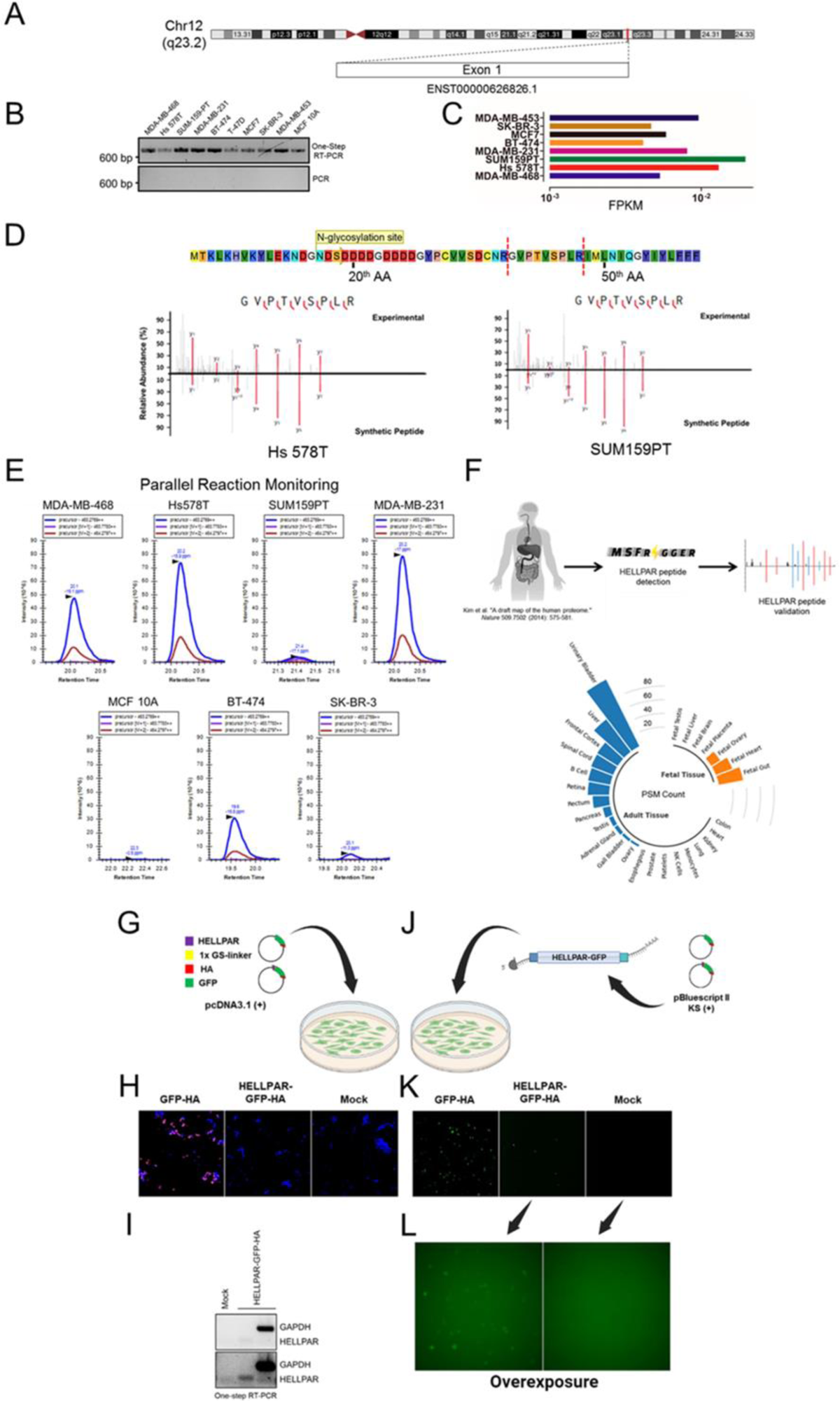
Detection, validation, quantification and expression of the protein encoded by the lncRNA-HELLPAR. (**A**) Genomic location and transcript architecture of the lncRNA-HELLPAR. (**B**) Transcript validation for lncRNA-HELLPAR across a panel of breast cancer cell lines using one-step RT-PCR and (**C**) RNA-seq. (**D**) Synthetic peptide validation of lncRNA-HELLPAR-encoded protein in Hs 578T and SUM159PT cells. Red lines represent y-ions. Briefly, cell pellets from Hs 578T and SUM159PT cells were processed as per Supplementary figure 2A and the identified peptide spectra (y-ions) for experimental/synthetic peptides were compared to determine similarity. (**E**) Parallel reaction monitoring of lncRNA-HELLPAR encoded protein peptide in various breast cancer cell lines. (**F**) Discovery of lncRNA-HELLPAR encoded protein peptide in publicly available human-derived adult and fetal tissues. Proteomics .RAW files obtained from Kim et. al., and lncRNA HELLPAR-encoded protein peptide spectral match counts were determined and plotted. (**G**) pcDNA3.1(+) constructs containing the lncRNA HELLPAR-encoded protein ORF tagged with C-terminal GPF-HA (lacking a start codon), or GFP-HA ORF were transfected into SUM159PT cells. (**H**) HA-tag protein expression was detected by immunofluorescence assay. (**I**) LncRNA-HELLPAR-GPF-HA tagged transcript was detected by one-step RT-PCR with primer probes targeting GFP. (**J**) pBlueScript II KS (+) constructs containing the lncRNA-HELLPAR ORF tagged with C-terminal GPF-HA (lacking a start codon), or GFP-HA ORF were in vitro transcribed, and the mRNAs were transfected into SUM159PT breast cancer cells. (**K**) GFP was detected by fluorescence microscopy 24 h post-transfection. (**L**) Fluorescence gain intensity was increased to detect low signal from lncRNA-HELLPAR-GPF-HA protein.

**Supplementary figure 4.**
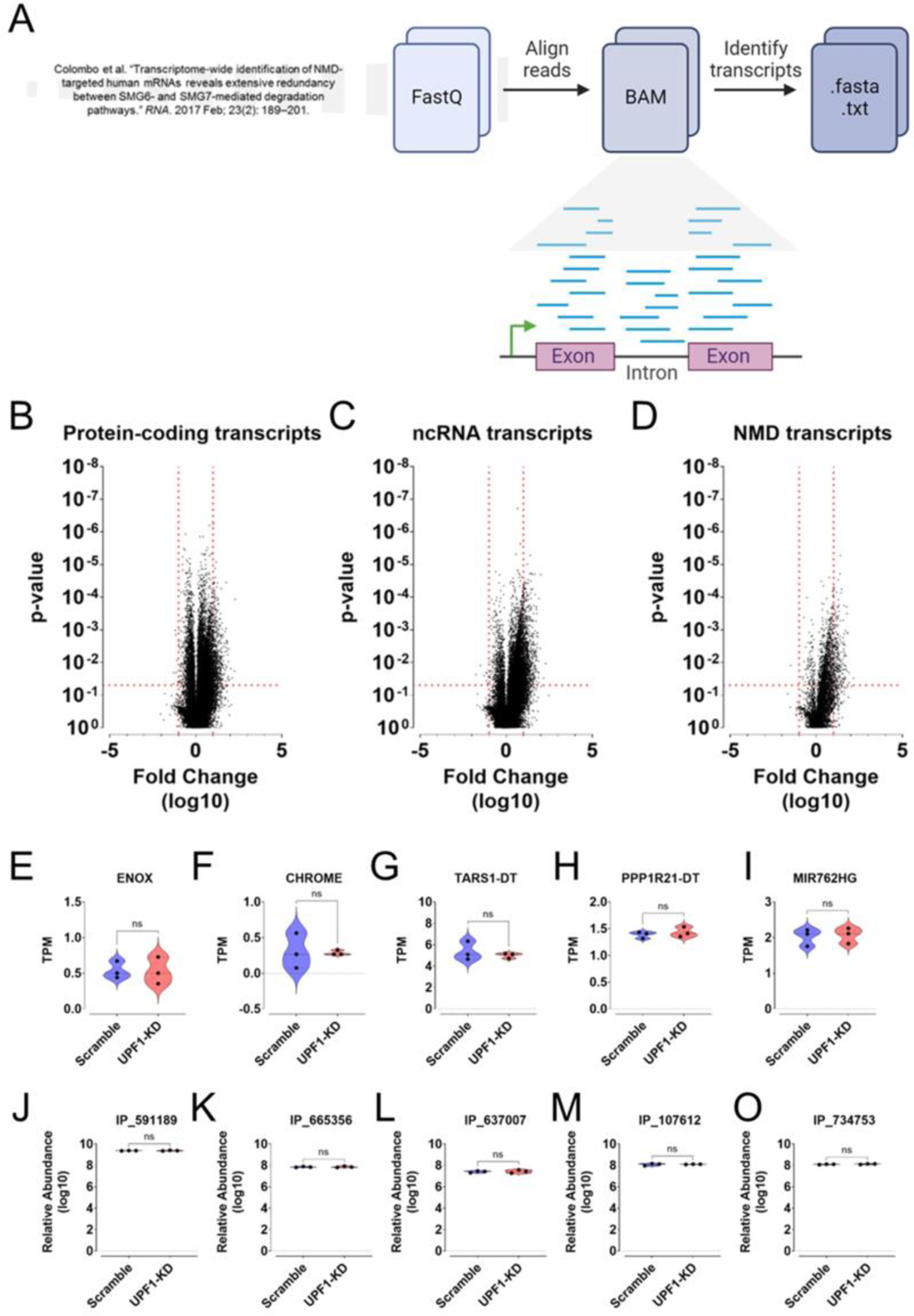
Nonsense-mediated decay targets the majority of ncRNA for degradation. (**A**) Workflow of data re-analysis of shRNA-mediated UPF1 knockdown in HeLa cells. Fastq files were pre-processed and re-analyzed using Salmon Quant. (**B – D**) Differential expression plot for (**B**) protein-coding, (**C**) ncRNA, and (**D**) NMD transcripts derived from HeLa cells after shRNA-mediated UPF1 knockdown. Log10 fold change values of transcripts were plotted; corresponding p-values were determined using two tailed t-tests. (**E – I**) SUM159PT and MDA-MB-231 cells were processed as per Supplementary figure 1A (transcriptomics). Detected transcripts were filtered for ncRNAs; well-characterized representative ncRNAs that were not modulated by UPF1 knockdown are shown. (**J – O**) SUM159PT and MDA-MB-231 cells were processed as per Supplementary figure 2A (proteomics). Representative novel proteins that were not modulated by UPF1 knockdown are shown.

