## Supplementary_Materials for "Regulation of Non-Canonical Proteins Encoded by Small Open Reading Frames via the Nonsense-Mediated Decay Pathway"

The raw proteomics files, along with their corresponding .pepXML files and the .params files of the proteomics search engine used, can be obtained through ProteomeXchange with the identifier PXD041585. To access these files, please log in with the following credentials: username - and password - 4ogvduJP. The raw RNA-seq files can be obtained through: https://dataview.ncbi.nlm.nih.gov/object/PRJNA956674?reviewer=fq8hhqddgcelvn8go92mqob19m

*Supplementary file 1:* Includes MultiQCs, HISAT-assembled transcripts, and TPM values in various formats for breast cancer cell lines within the 'BRCA_Panel_RNA-Seq' folder. Also included are unedited SDS-PAGE gels, protein length distribution, and normalized abundance values, among others, for low molecular weight protein enrichment QCs in the 'BRCA_Panel_Low_Molecular_Weight_Protein_Enrichment_QCs' folder.

*Supplementary file 2:* Includes a breast cancer cell line transcriptome-specific protein database, FragPipeGUI parameters, a list of identified proteins, heatmap PSM values, IonQuant MSstats converted values, and MSstat analysis of differentially modulated proteins within the 'BRCA_Panel_Novel_Known_Proteome' folder. Also included are files validating the novel protein HELLPAR, lncRNA-HELLPAR transcript FPKM values, identified peptide and synthetic peptide spectrum values, and PRM monitoring files, all housed in the 'HELLPAR_Novel_Protein_Validations' folder.

*Supplementary file 3*: Contains unedited western blot images, Salmon Quant output, TPM values, FragPipeGUI LFQMRB parameters, and a transcriptome-specific protein database in the 'UPF1_siRNA_Knockdown' folder. Re-analysis of PMID-27864472 RNA-seq data using Salmon is provided in the 'Re-analysis_Public_Data' folder. All supplementary files are accessible via Google Drive: https://drive.google.com/file/d/1I4LSy11WksgVzxBd8rXJTuf42Nk-HsDr/view?usp=drive_link
